# Prediction mismatch responses arise as corrections of a predictive spiking code

**DOI:** 10.1101/2023.11.16.567335

**Authors:** Kjartan van Driel, Lucas Rudelt, Viola Priesemann, Fabian A. Mikulasch

## Abstract

Prediction mismatch responses in cortex seem to signal the difference between an internal model of the animal and sensory observations. Often these responses are interpreted as evidence for the existence of error neurons, which guide inference in models of hierarchical predictive coding. Here we show that prediction mismatch responses also arise naturally in a spiking encoding of sensory signals, where spikes predict the future signal. In this model, the predictive representation has to be corrected when a mispredicted stimulus appears, which requires additional neural activity. This adaptive correction could explain why mismatch response latency can vary with mismatch detection difficulty, as the network gathers sensory evidence before committing to a correction. Prediction mismatch responses thus might not reflect the computation of errors per se, but rather the reorganization of the neural code when new information is incorporated.

## 1 Introduction

Strong neural responses in cortex to unexpected events are a common observation. The first well-known finding in this direction was that of mismatch negativity (MMN) in Electroencephalography recordings, where oddball stimuli result in elevated activity [1]. A long standing question is if MMN is a result of (bottom-up) neural adaptation or (top-down) prediction processes, and both possibilities have found experimental support [2, 3, 4, 5]. To separate these two effects, subsequent experiments went beyond the oddball paradigm and employed stimuli that become predictable only given a wider context. These experiments showed that also without stimulus adaptation neurons can show strong prediction mismatch responses (PMRs) [6, 7, 8, 9, 10, 11, 12]. While it is still a matter of ongoing research how exactly PMRs arise, they often seem to be modulated by top-down inputs to the cortical circuits [4, 13, 14, 15]. Together, these results suggest that top-down mediated predictions play a central role in cortical processing, and that PMRs are an important characteristic of their effect on neural dynamics.

What could be the computational function that underlies PMRs? The perhaps most discussed answer is given by classical hierarchical Predictive Coding (hPC) theories of cortical processing, which propose that neurons in cortex perform inference in a hierarchical model of sensory data [16]. Classical hPC argues that PMRs are generated by dedicated error neurons, which enable inference and learning by comparing top-down predictions to sensory observations [17], and thus it connects the observed top-down modulated PMRs to a specific cortical function. Following this theory, several models of spiking neural networks have demonstrated possible mechanisms that could lead to the presence of error neurons in cortical circuits [18, 19, 20, 21] (or combined prediction and error neurons [22]). However, while these models give mechanistic accounts of PMRs, so far there is no direct connection of spiking neuron PMRs to the formal model of inference that is at the heart of the predictive coding theory [23]. This issue is connected to the more general open question of how the inference and learning algorithm of classical hPC might be implemented by spiking neurons [23, 24].

In this paper we offer an alternative account of PMRs, based on theories of hierarchical inference in cortex that operate without error neurons [24] (see also [25, 26, 27, 28]). In summary, we assume that the neural spiking code in cortex is predictive, and a mispredicted stimulus requires a correction of the code, which leads to additional neural activity. This will allow us to connect PMRs to biologically plausible theories of inference and learning in spiking pyramidal cells [24, 29]. Finally, we will discuss how PMRs might provide an opportunity to experimentally distinguish between models of hierarchical inference with or without error neurons.

## 2 Theory

To illustrate our ideas with a particular example, our aim is to model an experiment by Fiser and colleagues [9]. In this experiment a mouse traverses a virtual tunnel with landmarks (i.e., the location in the tunnel is known to the animal). At specific locations the mouse is then presented with visual patterns that are either predicted by the location, or mispredicted (Fig 1a). In the experiment, mispredicted stimuli lead to heightened neural activity in visual cortex V1 [9].

**Fig 1.**
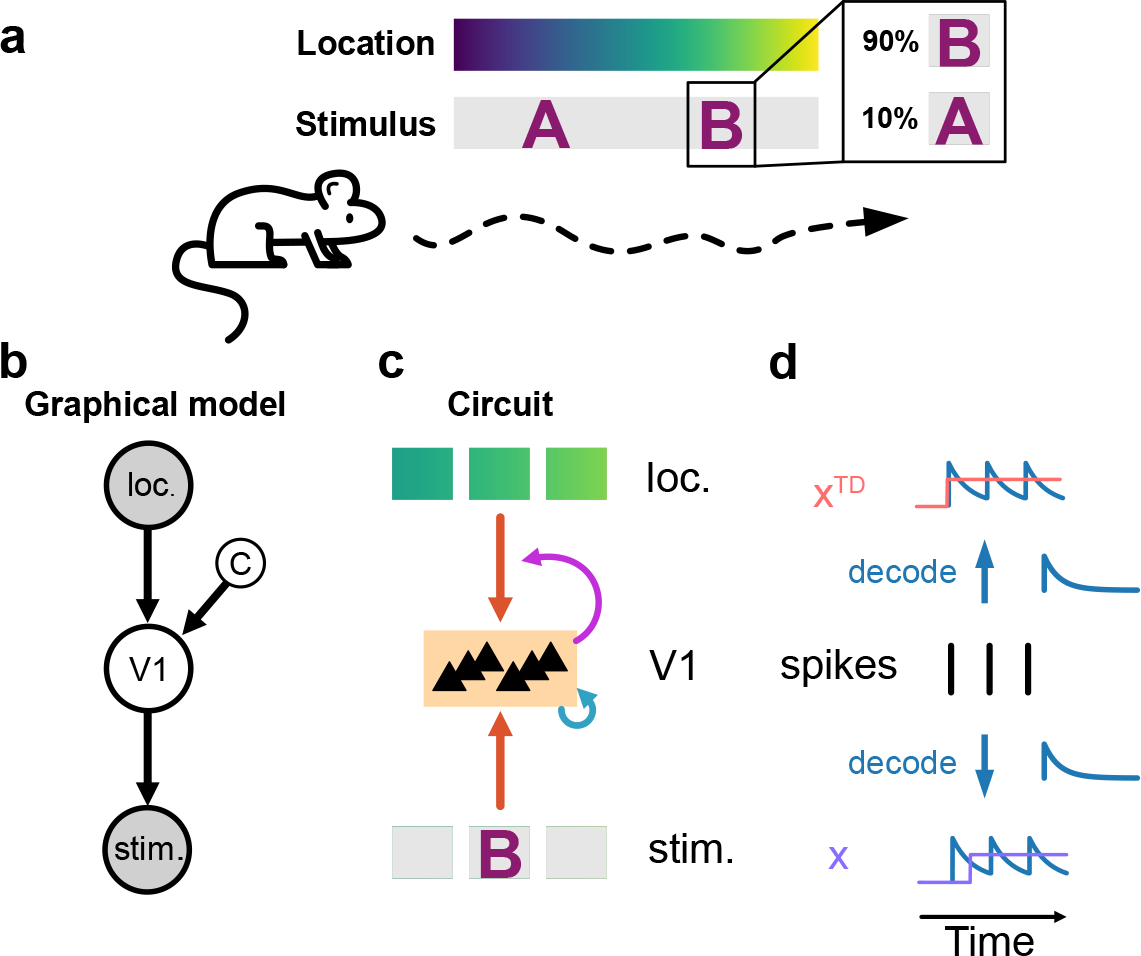
Model of neural coding for spatially predicted visual stimuli. **(a)** Neurons aim to encode visual stimuli (A,B) that are associated with specific given locations of the animal, as in experiment [9]. In rare cases the association can be violated, i.e., 10% of the trials the predicted stimulus B is replaced by A. **(b)** We assume the animal forms a generative model of visual data. The location (loc.) predicts the stimulus representation in V1, and the stimulus representation predicts the sensory observation (stim.). An additional context variable (*c*) indicates if the prediction from location is valid. Shaded grey circles denote signals that are given (fixed) by the experiment. Variables in white circles have to be inferred to model the given data. **(c)** Cortical circuits invert the generative model. Depending on the inferred context variable *c* the influence of top-down location information is enabled or disabled. **(d)** Illustration of how spikes in V1 aim to track visual (*x*) and location (*x*^*T D*^) input signals. Spikes encode the future signals via an exponential kernel, or in other words, they predict the future signal. Spikes are fired such that they simultaneously conform to visual and location signals.

The basic assumption of our model is that V1 aims to find a predictive spike encoding of visual stimuli. This assumption has two motivations: i) in experiment neurons have been found that learn to predict upcoming stimuli [9, 30], and ii) it might be advantageous to predict stimuli to facilitate robust and rapid stimulus recognition [30, 31, 32, 33]. We realize this idea formally, by assuming that the mouse has an internal (generative) model of how sensory stimuli are generated (Fig 1b,d), which is inverted by the neural circuits in cortex (Fig 1c). More specifically, the generative model states that the location is predictive of the encoding in V1, and the encoding is predictive of the perceived stimulus. Therefore, to find the encoding in V1, neurons have to combine sensory signals and location information (e.g., provided by the anterior cingulate cortex [9]).

An important additional component of the generative model is a binary context variable *c* ∈ {0, 1}, which determines if the location is indeed predictive for the encoding (*c* = 1) or not (*c* = 0). This context variable is necessary to enable the generative model to capture the sensory data in cases where the location mispredicts the observed stimulus. Intuitively, this enables the mouse to realize that a predicted stimulus (e.g., an object) is not present at a certain location, and to integrate this information into the stimulus representation. Otherwise, the mouse would continue to naively combine the wrong prediction from location with the sensory observation, resulting in a wrongfully biased representation no matter how much evidence for the inadequacy of the prediction is available. Finding the context variable while inverting the generative model is a nontrivial problem [34, 35]. To tackle this, we here build on previous work [34] and deterministically switch from *c* = 1 to *c* = 0 if the location prediction deviates significantly from the encoding in V1 over an extended period, which can be motivated from the generative model (i.e., this implements a maximum a-posteriori estimate of *c*, see Methods). In the neural circuit, the realization that the location is mispredictive (*c* = 0) then leads neurons in V1 to ignore inputs from location neurons (Fig 1c). We will outline possible biological mechanisms for this computation in the Discussion.

## 3 Results

To demonstrate how this model leads to heightened neural activity during prediction mismatch, we simulated the responses of a small network to simple experimental stimuli. A network of 6 neurons encodes two patterns (A and B) and an inter-stimulus signal, denoted by low-dimensional orthogonal vectors. For simplicity, stimulus encoding weights and top-down connections were set fixed. Top-down weights were set such that a location predicts the presence of a specific pattern in the network, which reflects how the patterns are presented in experiment (Fig 1a). In principle these weights could also be learned via voltage-dependent plasticity rules [24, 29, 36, 37].

We now explain how a prediction mismatch leads to a correction of the population code, and with that to a burst of neural activity. Because a location is predictive, neurons coding for the predicted pattern (B) will be driven by top-down location inputs, and fire in anticipation to encode the future signal (Fig 2). When the actual pattern (A) appears, the network starts to find an encoding of the stimulus (A) that is biased towards the prediction (B). The emerging mismatch between prediction and encoding leads the network to realize that the prediction is invalid (i.e., *c* switches from 1 to 0). In consequence, the top-down prediction is ignored and the encoding in the population is corrected from the predicted pattern (B) to the observed (A). In our model, this correction requires activity of two types of neurons: i) neurons coding for pattern A, which are driven by bottom-up input, and ii) neurons coding for pattern ‘-B’, which are driven by neurons coding for B and released from top-down inhibition (Fig 2b, Suppl. Fig S1). These ‘correction neurons’ (ii) are required because past spikes predicted pattern B, and this prediction has to be removed from the predictive encoding. Intuitively, a predictive code has ‘momentum’, and a rapid correction requires strong network action in the form of spikes.

**Fig 2.**
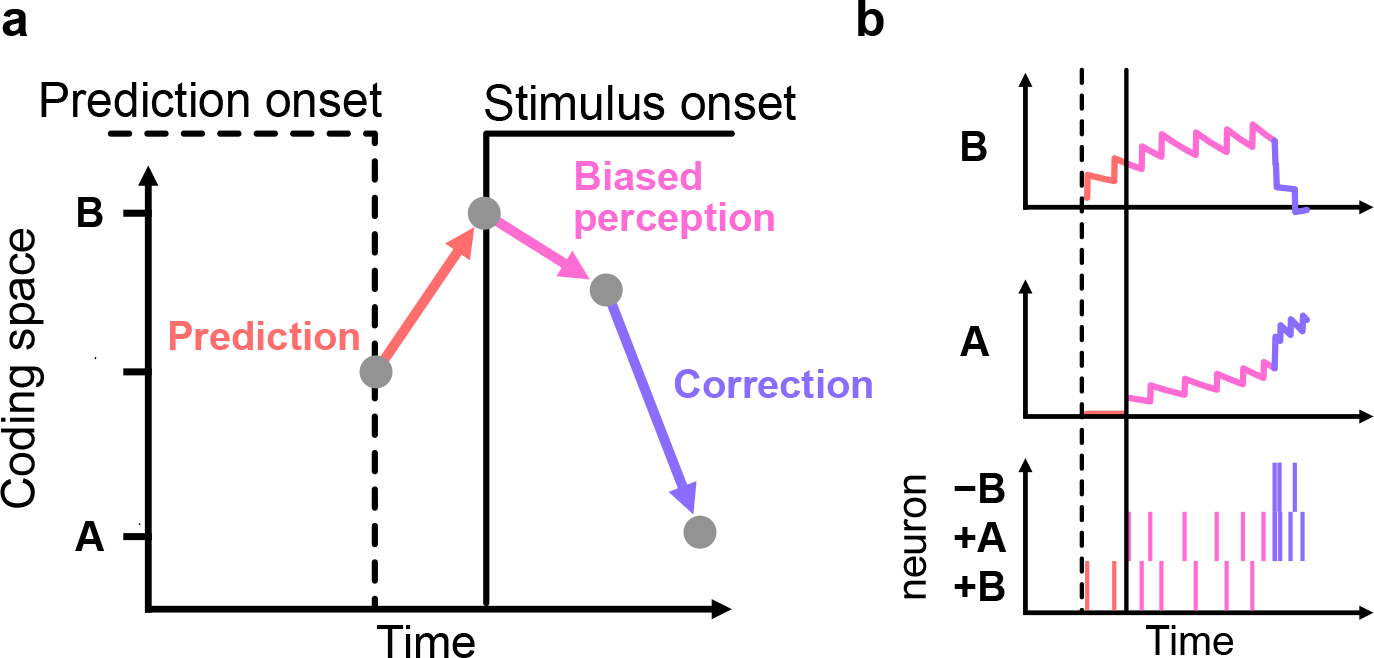
A mechanism for PMRs through corrections of a predictive spiking code. **(a)** Schematic illustration of the evolution of the population code over time in case of a prediction mismatch. After prediction onset the population begins coding for the predicted pattern (B). With stimulus onset the network starts to encode the observed pattern biased towards the prediction. When the prediction mismatch is detected (*c* switches from 1 to 0), the code is corrected by adding the observed pattern and removing the wrongly predicted pattern. Moving in coding space (arrows) requires neural spiking. **(b)** The same evolution of the population code but in simulation. Top panels show the decoded stimulus code, bottom panel shows network spiking activity. Because past spikes predict the future signal, a rapid correction requires additional activity: i) to encode the correct pattern, here via a burst of activity in stimulus coding neurons (+A); ii) to remove the population code for the predicted pattern, here via dedicated ‘correction neurons’ (-B).

In our simulations, these two processes together resulted in a burst of activity of the population in response to mispredicted stimuli (Fig 3a). This increase in activity was even more pronounced for correction neurons (‘-B’ neurons), which only become active when the population code over-predicts a pattern (B) (Fig 3b). Since correction neurons remove wrong predictions from the population code, they become more active in trials where the activity of mispredictive neurons before the stimulus was stronger (Fig 3c), which was also found experimentally [9]. In that sense, these neurons can be considered dedicated ‘mismatch’ or ‘error’ neurons, although they simply keep the population code in check by removing the over-predicted pattern from the code. Note that, in principle, the increase in activity could also arise without correction neurons, but the resulting encoding would be slower to capture the observed stimulus correctly.

**Fig 3.**
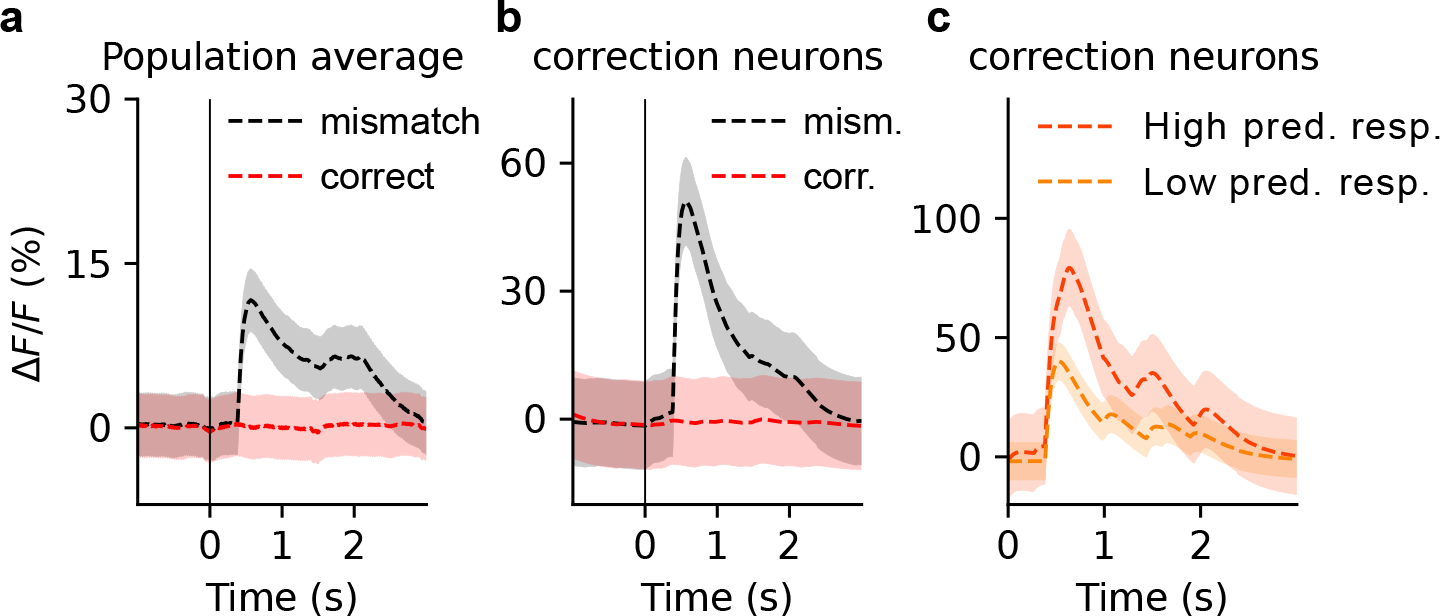
The model recovers several effects of PMRs as observed in experiment [9]. All panels show simulated fluorescence recordings based on simulated spike trains (see Methods). Stimulus onset is at Time = 0s. **(a)** The average population response to a mispredicted pattern (A) is higher than to a correctly predicted pattern (B). **(b)** This effect is even more pronounced in neurons that remove the over-prediction of the mispredicted pattern (‘-B’ neurons), and therefore appear as dedicated ‘correction neurons’. **(c)** Same as the mismatch condition in panel b, but with trials separated based on the strength of predictive activity that precedes stimulus onset. Stronger predictive activity requires a stronger response of correction neurons. Note that in the modeled experiment these effects (**a** - **c**) are found for the omission of a predicted stimulus (Fig 4 in [9]). To ease interpretation we here consider a misprediction (B instead of A), but in our model this is equivalent to a omission (B instead of inter-stimulus signal). Panels **a** and **b** show mean and standard deviation for 50 trials in each condition. Panel **c** is based on the same data and shows top and bottom 10% of trials sorted by pre-stimulus activity.

Finally, we aimed to analyze the proposed context detection mechanism (switching of variable *c*) in more detail. *c* was estimated by selecting the context (‘correct prediction’ or ‘misprediction’) which better captures the relation of encoding and prediction. This was implemented by comparing the time-window averaged prediction error (context criterion) to a switching threshold (Fig 4c). We found that this implies a longer time delay before a context switch if the mismatch is harder to detect (e.g., if it is smaller), which we verified in our simulations (Fig 4). Intuitively, in situations where it is hard to detect the inadequacy of the prediction the mouse has to deliberate longer, and gather more evidence, before the internal predictions can be ignored.

**Fig 4.**
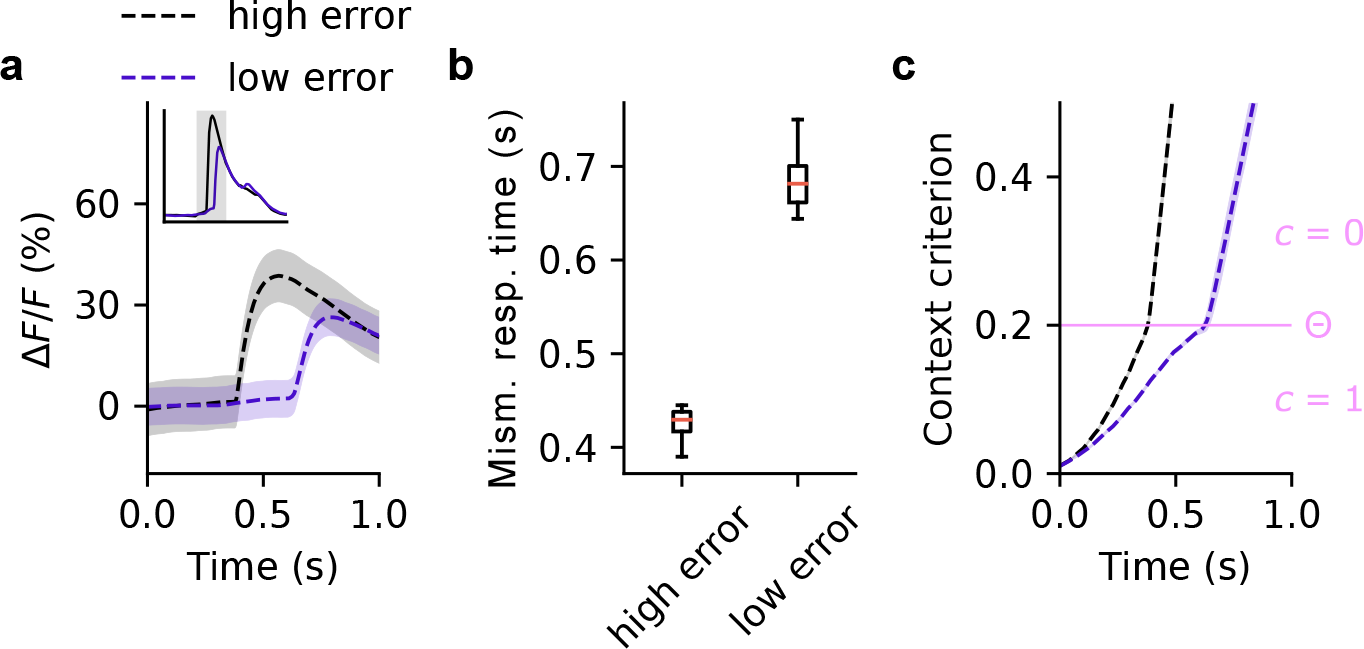
Mismatch detection difficulty influences latency of model PMRs. To show this the network was presented with a pattern that is a mix of the predicted (B, 30%) and the unpredicted pattern (A, 70%), which means the network observes only a partial mismatch between prediction and stimulus. **(a)**If the mismatch of prediction and observation is smaller (low error), a longer latency of PMRs can be observed in the response of correction neurons (i.e., -B neurons). Inset shows the same responses over a longer time-window. **(b)** The median delay of the mismatch response (here simply measured as the time of Δ*F/F* crossing 20%) is several hundred milliseconds longer. Note, that this constitutes only a qualitative prediction of our model, and smaller or larger delays could be obtained by choosing different parameters (e.g., Θ). **(c)** The difference in latency originates from the dynamics of the criterion for switching the context variable *c* (time averaged prediction error), which increases with a slower rate in case of a partial mismatch. This results in the network reaching the threshold for initiating a correction of the encoding (Θ) later. All panels show results for 50 trials in each condition. Outliers are not shown in panel **b**.

## 4 Discussion

Building on previous work on efficient spike coding [24, 38], we here proposed that prediction mismatch responses (PMRs) can result from the correction of a predictive spiking code. In our model, the correction is initiated when the prediction from other areas is incompatible with the activity in the coding population over an extended time. When this happens, the old prediction is removed from the population code and the representation of the perceived stimulus is added, both of which requires additional neural activity (Fig 2). These dynamics are consistent with experimentally measured responses of V1 neurons during prediction mismatch (Fig 3).

The correction dynamics were derived from a generative model view of perception (Fig 1). Specifically, we assumed that the internal model of the mouse distinguishes between two cases: One, where the location is predictive of the encountered pattern, and one where it is not. Which one of these two possibilities is the case has to be inferred from observations, and we showed that this process takes longer in cases where the observations are more ambiguous (Fig 4). As an illustrative example, consider that you have left an object (e.g., a bottle) in a room, and another person removed it later. Entering the room you have the expectation to observe the bottle (and are more inclined to perceive it), but after a short look you realize that the location-object prediction you held was incorrect and you correct your internal model. If, instead, the room would be only dimly lit, it would take longer for this realization to occur, and other bottle-shaped objects might deceive your perception in the meantime. Based on our model, we argue that this realization and the subsequent correction of the representation of sensory stimuli is what underlies the PMRs observed in experiment.

## Experimental predictions

Our model makes three key predictions:

i. When neurons in a population show dedicated mismatch responses resulting from the proposed mechanism, the same population will also contain stimulus predictive neurons (or, at least, stimulus selective neurons that are strongly biased towards the prediction). In our model these neurons cooperate to implement an efficient and responsive spiking code [39]. A possibly similar co-location of mismatch- and stimulus-selective neurons has been found in several experiments [9, 13, 40].
ii. Our model predicts that mismatch responses emerge with a longer delay when detecting the mismatch becomes more difficult (e.g., for a smaller mismatch), since more evidence has to be gathered before committing to a correction (Fig 4). This could be tested by measuring the time to the onset of the mismatch response depending on the stimulus noise level, or mismatch size. There have been several experiments showing the predicted (or a similar) effect in EEG recordings of MMN [5, 41, 42, 43, 44, 45, 46]. We expect that this effect can also be measured in single neuron recordings of PMRs, where especially mismatch-selective neurons should be affected.
iii. Our model predicts distinct origins for the driving connections in positive and negative PMRs (Supplementary Fig S1). Positive PMRs (i.e., the stimulus is bigger than expected) result from excess drive through bottom-up connections, which is not cancelled by lateral inhibition in the population. Negative PMRs, in turn, result from drive within the population, which is not cancelled by top-down inhibition (these neurons are the ‘negative’ coding neurons that appear as dedicated error neurons; Fig 3). In practice, however, it might be difficult to achieve situations where these two types of PMRs can be observed in isolation, and in our simulation both cases appear. Note, that in both cases the PMRs are nevertheless a result of mispredictions, typically from top-down inputs. Top-down inputs in our model, however, do not directly drive the delayed PMRs, but only indirectly cause them when the wrong predictions they have provided are finally ignored.

### How does this model differ from other explanations of mismatch responses?

Previously, prediction mismatch responses have been explained with the dedicated error neurons that are proposed by classical hPC and similar models [17, 47]. While some neurons in our model show responses that appear as dedicated error responses (Fig 3), the interpretation of the purpose of PMRs in our model and in classical hPC is very different. In classical hPC, the responses of error neurons can only be interpreted in conjunction with associated activity in prediction neurons that are updated on error activity [16]. In our model, mismatch responses simply constitute a part of a predictive population code that is meaningful without any additional information. In both interpretations, mismatch responses can be understood to signal that the prediction that was conveyed up until that point was wrong, and that any higher-level area might have to adapt to this (e.g., by changing the planned course of action in motor ares). In contrast to classical hPC, however, our model does not require feedback-signals from every targeted higher-level area that cancel the prediction error.

Experimentally, classical hPC and our model might be distinguished in two ways. First, by looking at the inputs that drive mismatch responses. In error neurons, PMRs are typically assumed to arise when bottom-up drive and top-down inhibition do not match (Supplementary Fig S1), as opposed to the different origins in our model we have discussed before. Second, by looking at the temporal dynamics of PMRs. As we have discussed, our model predicts that the latency of PMRs depend on the difficulty of detecting a mismatch, e.g., the size of mismatch (Fig 4). In contrast, in error neurons the size of mismatch would not be expected to have an impact on the latency, since here neurons are expected to respond as early as possible [18, 19, 20] (Note, that this is different for noise in the stimulus, which might delay the response of precision weighted error neurons [48]). This means that the same experiment we have proposed to test our model in the last section might also be used to distinguish between theories with or without error neurons.

Another set of work has proposed that prediction mismatch responses could be a signature of an efficient adaptive code [34, 35]. In this idea the neural code is continually adapted to most efficiently encode a signal, and a temporary mis-adaptation results in an inefficient encoding, that is, increased activity. This means that in this theory, similar to error neurons, the mismatch response is immediate, and ceases with the adaptation. We have employed the same idea of an adaptive code, but showed that in a population code the correction *after* an adaptation causes a surge of activity. Therefore, our model constitutes an extension of these adaptive coding models and is compatible with their ideas. Future models might look more in detail at the possible interaction of these two proposed components of mismatch responses that arise early and late after stimulus onset.

Yet another explanation for PMRs is based on the idea of neural sampling [49]. Here, a prediction mismatch is thought to increase the uncertainty (variance) of the inferred posterior over latent variables, which in turn results in increased neural activity (owing to the non-linearity in neural spiking) [26]. We find this effect also in our model, as after detecting a mismatch the effective variance of the inferred posterior is increased by ignoring top-down predictions. This results in an overall increased population activity even after the initial correction response (Fig 3a). However, it is not clear how this alone could lead to mismatch selective neurons (Fig 3b) and mismatch size dependent PMRs (Fig 3c) as observed in experiment [9]. Thus, it is possible that the marked rise in activity during PMRs is a result of both, a correction response and an increased variance in neural sampling.

In previous work we have also proposed a model for the emergence of multimodal mismatch responses [50]. In this model, mismatch responses arise when different areas simultaneously encode sensory information, as these areas compete to encode the signal and thereby cancel activity in their respective partner area. Here the proposed purpose of mismatch responses (i.e., encoding the signal that is not explained by the other area) is very different from the presented model, but there is no reason why not both of these computations could exist alongside each other. Based on our models we argue that different types of mismatch responses in cortex can have strictly different meanings in cortical computations, and it might not be appropriate to describe all these observations with a single computational principle.

### Limitations

In a similar vein, the assumptions of our model might not be appropriate for all model corrections an animal can perform. We here assumed that the top-down predictive signal (i.e., the location) is perfectly certain and fixed, which makes sense in the modeled experiment where the mouse can orient itself with landmarks [9]. However, if the animal is not certain of the content of higher-level representations, in light of new sensory evidence these higher-level representations might be corrected (instead of the ‘coupling’ between levels, as in the presented model). This type of correction likely has somewhat different temporal dynamics than the one presented here, and future work could investigate these dynamics in a more complex multi-level model.

Overall our model was designed as simple as possible to give an intuitive understanding of how PMRs can arise in spiking neurons, but might lack features that determine how these responses realize in the brain in detail. Most crucially, all neural connections in this model have been fixed to perform inference in a fixed generative model (Fig 1), but in general the generative model and the associated weights would have to adapt to the stimulus statistics. Doing this would lead top-down weights to learn predictive connections between location and stimulus [24, 29], and bottom-up weights to learn efficient stimulus representations [36, 37]. Especially how stimuli are represented (which includes the tuning of ‘correction neurons’; Fig 2) can have important implications on how exactly the resulting PMRs look like. For example, in some experiments [51] PMRs seem to be less specific than suggested here, which might be a result of the circuit encompassing less precisely tuned ‘correction neurons’ than we have assumed here.

It is also possible that additional mechanisms, which have not been modeled here, play important roles in the generation of PMRs. In particular, attention to interesting (in this case, unexpected) stimuli could enhance neural representations and thus introduce increased activity [26, 52]. A challenge with this idea is that there seem to exist several different forms of attention [52], and PMRs can occur independently of at least some of them [2, 5, 53]. Nevertheless, the presence of such mechanism of activity enhancement, which might occur in parallel to the mechanism proposed here (possibly based on the same deviance detection mechanism), could alter how neural activity changes during PMRs in detail. Alternatively, attentional enhancement could also be the main driver of PMRs and it would be important to formulate such models in more detail, in order to distinguish them from the presented correction based account of PMRs.

### Implementation in cortical circuits

So far we have operated with an abstract model, but in the context of a theory of dendritic predictive coding in cortex [24] we can speculate about possible biological implementations of the proposed mechanism. Consistent with the proposed model, this existing theory proposes that layer 2/3 pyramidal cells find a predictive encoding of sensory stimuli, where bottom-up sensory information arrives at basal dendrites, and top-down predictions from other areas at apical dendrites. Thus, the only component in our model that so far has no biological interpretation is the adaptive switch that allows/disallows top-down predictions to influence the population code (i.e., variable *c*; Fig 1). Since top-down predictions arrive at apical dendrites, one possibility would be that the coupling of the apical dendrite to soma is adapted [54], which would imply that apical dendrites decouple from the soma during a mismatch event. Similar mechanisms have been proposed to be responsible for adaptive associations in cortex [55] or the conscious processing of sensory stimuli [56]. Another, less intrusive mechanism would be that apical inhibition precisely controls the impact of specific predictions on neural activity by cancelling apical inputs in case of a prediction mismatch. Indeed, somatostatin-expressing (SST) interneurons, mostly targeting the apical dendrite, seem to play a central role in the generation of PMRs [4, 51, 57]. This proposal leads to the seemingly paradoxical prediction that apical inhibition of pyramidal cells through SST interneurons would be decreased when top-down inputs are predictive, but increased when they are mis-predictive and pyramidal cells show strong mismatch responses (when otherwise SST and pyramidal neuron activities tend to be anti-correlated [58]), for which there are some indications [57].

While these biological connections are speculative, they provide testable predictions for how exactly cortex can adapt its internal model in cases where internally generated predictions do not match sensory observations. Testing for the biological mechanism behind these PMRs might be important in order to understand the causes of mental disorders where PMRs are altered, such as schizophrenia or certain learning disorders [59].

## Acknowledgements

We want to thank Anna Vasilevskaya, Fabian Sinz, Georg Keller, Suhas Shrinivasan and the Priesemann Lab, especially Andreas Schneider, for helpful discussions. F.A.M. and L.R. were funded by the German Research Foundation (DFG), SFB1286. V.P. received support from the SFB1528, Cognition of Interaction. V.P. and L.R. were supported by the MBExC Excellence Cluster, Deutsche Forschungsgemeinschaft (DFG) under Germany’s Excellence Strategy - EXC 2067/1- 390729940.

## Author contributions

**KvD**: Investigation, Software, Writing - Original Draft. **LR**: Investigation, Writing - Review & Editing. **VP**: Supervision, Writing - Review & Editing. **FAM**: Conceptualization, Investigation, Supervision, Writing - Original Draft.

## Methods

### Generation of input signals

We model the experimental setup of Fiser and Keller [9] by defining the input signals model V1 receives. Visual inputs **x**^∗^(*t*) were 2 different patterns (A and B) and an inter-stimulus signal, which were presented with a one-hot encoding. The location in the tunnel 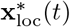 was assumed to be perfectly known and similarly given by a one hot encoding akin to the representation in place cells, which each corresponded to the location of a pattern (or inter-stimulus signal) in the tunnel.

Patterns were presented for 1.5 s before switching to the next pattern. Input signals were low-pass filtered to simulate the integration in visual cortex 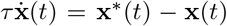. The location signal was filtered similarly, but presented shortly (100 ms) before the stimulus to achieve anticipatory spiking. Intuitively, we argue that the mouse reaches a location in the tunnel where it predicts the pattern shortly before it actually observes it. Note, that for the effect of mismatch responses this is not strictly necessary, but a choice we made to model this specific experiment.

### Generative model of sensory data

To simulate the perception process of the mouse, we first defined a generative model we assume the mouse has of sensory data, and then found a network that sampled from the inverted model (i.e., the posterior for the variables of interest).

The model was defined via a hierarchy of Gaussian distributions, where stimuli **x** were generated by activity *V* 1 corresponding to the hidden causes of sensory data (here the identity of the pattern)

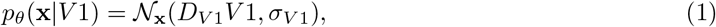

and hidden causes *V* 1 were generated by the location of the mouse **x**_loc_

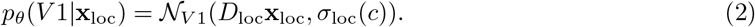

For simplicity, we set all parameters *θ* = {*D*_*V* 1_, *σ*_*V* 1_, *D*_loc_, *σ*_loc_(*c*)} by hand. The decoder weights *D*_*V* 1_ and *D*_loc_ might in principle be learned using voltage-based plasticity rules [24, 29], but were here chosen as the identity matrix.

To model the fact that objects (i.e., patterns) can be absent from a certain location, the variance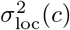 of the location prediction of *V* 1 is adaptive. Specifically, 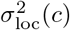 depends on a binary context variable *c* ∈ {0, 1} which indicates if the location is predictive (*c* = 1) or not (*c* = 0)

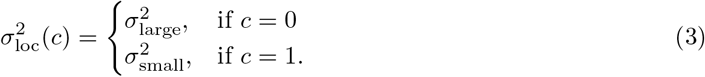

The (implicit) prior distribution *p*_*θ*_(*c*) we use for *c* will be discussed later. A similar adaptive variance has also been proposed in other models [60, 61], where, however, the variance is typically adapted continuously. In our scenario, in contrast, we assume that a binary variance captures better how the data is generated (i.e., the location is predictive or not, but nothing in between).

### Neural dynamics performing inference through sampling

The goal of neural inference is to find an approximation to the posterior *p*_*θ*_(*V* 1 | **x, x**_loc_) using a neural network. To this end, we replace the continuous variable *V* 1 by a spike based representation, similar to previous work on spike-based representations [36, 62]. We define *V* 1 = *D***r**(*t*) as a transformation of neural responses **r**(*t*), generated by 6 neurons. The transformation matrix *D* was defined as

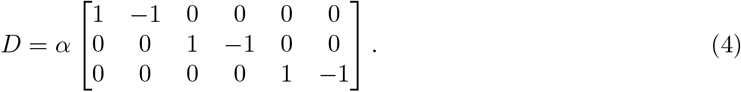

Neural responses are defined as a readout of neural spiking *s*_*i*_(*t*) ∈ {0, 1} convolved with exponential spike traces *r*_*i*_(*t*) =Σ _*t*_*′*_≤*t*_ *κ*(*t* − *t*^′^)*s*_*i*_(*t*^′^), where *κ*(Δ*t*) = exp(−Δ*t/τ*). Neural spikes are thus read out in the future, and can be considered to constitute a prediction of the upcoming signal.

For spike based inference, we introduce an additional prior on neural spiking

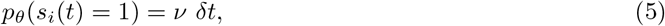

where *δt* denotes the physical time elapsed between successive timesteps. This prior can also be thought of as a metabolic cost on neural activity.

Representing *V* 1 in this manner, combined with the additional prior allows us to heuristically derive a network of stochastic spiking neurons performing inference. The derivation relies on two key observations. First, we can write the spiking probability for neuron *i* in terms of log-probabilities as

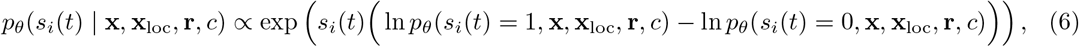

which is similar to previous approaches for spike-based neural sampling [63], and can be considered a stochastic generalization of the spike-by-spike framework [36]. We note that in general the spiking probability for neuron *i* is not independent from the rest of the network. Our second observation is that if we suppose *δt* to be small our metabolic prior in Eq. (5) forces the probability of simultaneous spiking to approach zero, and therefore introduces an independence between between spiking.

By using these observations and writing out the difference in logarithms in Eq. (6) for our choice of model and representation we find that the spiking probability of neuron *i* can be expressed as

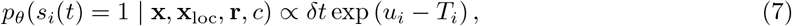

for a membrane potential *u*_*i*_ and threshold *T*_*i*_ given by

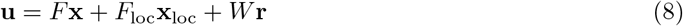

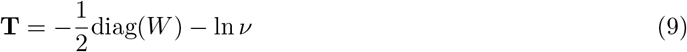

with

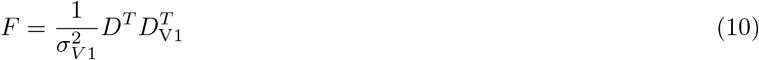

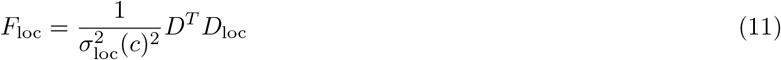

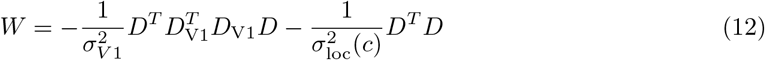

where diag(*W*) denotes the vector containing the diagonal elements of *W* and the superscript on *D*^*T*^ denotes the matrix transpose. This form of the spiking probability can be seen as a special case of the spike response model with exponential escape noise [64]. Alternatively, similar results could be obtained by simulating neural spiking as performing maximum a-posteriori inference in the generative model, which can be implemented with similar but deterministic neural dynamics [36]. To obtain fluctuation based results (e.g., Fig 3c), however, additional noise on neural inputs would be required in this case.

### Context switching algorithm

The context variable *c* that indicates the validity of the location prediction also has to be inferred. We performed this inference using an approach that has been proposed in previous research [34, 35]. At time *t* we selected *c* as the maximum a-posteriori estimate for the recent past time-window *T* : *c* ← arg max Σ_*t*_ −_*T<t*_*′*_*<t*_ log *p*_*θ*_(*c* | *V* 1(*t*^′^), **x**(*t*^′^), **x**_loc_(*t*^′^)). Since *c* is binary we selected *c* = 0 iff

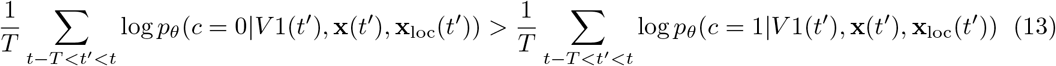

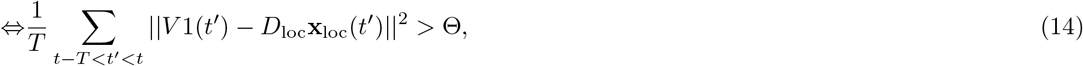

where Θ is a switching threshold that combines the prior on *c* and the variance of the model *p*_*θ*_(*V* 1 | **x**_loc_). For simplicity, instead of explicitly defining a prior on *c* we directly chose the switching threshold Θ.

### Simulation of calcium fluorescence signals

To simulate fluorescence signals Δ*F/F* as measured in experiment we first created calcium traces for each neuron by convolving the spike train with a realistic fluorescence kernel

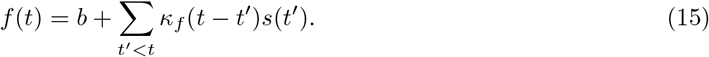

Here, *b* = 4.0 is a baseline activity in the signal which models average measurement noise and other activity, and re-scales the normalized signal Δ*F/F* . The kernel of the fluorescence elicited by a spike was defined by *κ*_*f*_ (Δ*t*) = exp(−Δ*t/τ*_decay_)(1 −exp(−Δ*t/τ*_rise_)), with *τ*_rise_ = 80ms and *τ*_decay_ = 400ms, as measured in experiment [65]. The normalized fluorescence signal was then computed via

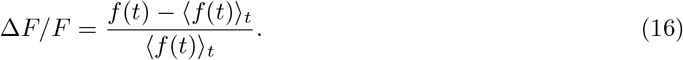

### Parameters

| Parameter | Value |
| --- | --- |
| $\delta t$ | 1 ms |
| $\tau$ | 200 ms |
| $\alpha$ | 0.15 |
| $\sigma_{V1}^2$ | 1/300 |
| $\sigma_{\text{small}}^2$ | 1/300 |
| $\sigma_{\text{large}}^2$ | 1 |
| $\nu$ | 13.60 Hz |
| (context criterion window) $T$ | 500 ms |
| (context criterion threshold) $\Theta$ | 0.2 |

### Simulation code

Code to reproduce the main results of this article can be found on https://github.com/Priesemann-Group/prediction_mismatch.

## Supplementary figures

**Fig S1.**
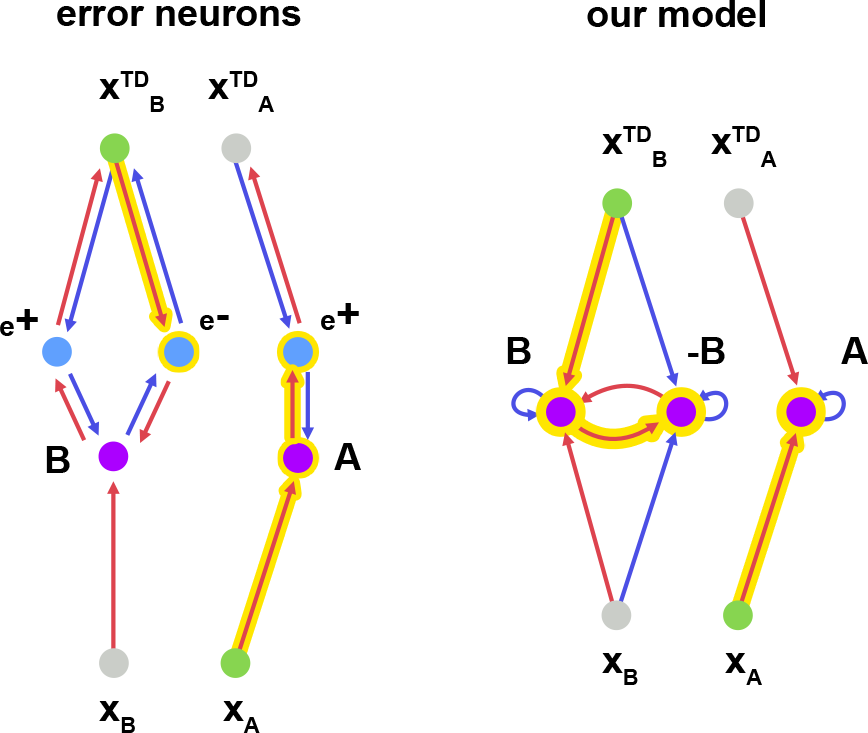
The driving connections of PMRs are slightly different in classical hPC and our model. Arrows denote excitatory (red) and inhibitory (blue) connections. Yellow arrows indicate the path of activity that leads to PMRs. Note, that in our model mismatch neurons (-B) only become active once top-down inhibition ceases after the correction is initiated. Note also, that in our model +B and -B neurons excite each other, but their effect on downstream neurons (e.g., in motor areas, not depicted) would be opposing. The mutual excitation ensures that downstream neurons receive the appropriate net-excitation: If one of the neurons was too active (e.g., through a misprediciton), it will excite the other (opposing) neuron in order to correct the net-excitation in downstream neurons.

